# Deep imaging revealed dynamics and signaling in one-to-one pollen tube guidance

**DOI:** 10.1101/2023.04.19.537439

**Authors:** Yoko Mizuta, Daigo Sakakibara, Shiori Nagahara, Ikuma Kaneshiro, Takuya T. Nagae, Daisuke Kurihara, Tetsuya Higashiyama

## Abstract

In angiosperms, pollen tube guidance allows sperm cell delivery to the female gametes within the ovule, which are deeply embedded in a flower. However, when an ovary includes multiple pollen tubes and ovules, it is unclear how each ovule is fertilized one-to-one by a pollen tube. Here, our two-photon imaging revealed the pollen tube dynamics in living ovaries. The number of pollen tubes and ovule maturity affected the target selection among multiple ovules. On the inner surface of the septum epidermis within the transmitting tract, pollen tube behavior and emergence were regulated by the ovular sporophytic signals. In funicular guidance, the second pollen tube was strictly repelled by the FERONIA and LORELEI-dependent gametophytic signal, especially more than 45 minutes after the first pollen tube had passed. Such highly spatiotemporal regulation mechanisms in the one-to-one pollen tube guidance may allow angiosperms to produce more offspring in nature.

---

Flowers are the reproductive organs of angiosperms that evolved between the Jurassic and Lower Cretaceous Periods^1^. Inside a flower, two sperm cells fuse with an egg and a central cell, respectively, initiating seed development. However, sperm cells have lost their motility to evolution and require the pollen tube to be delivered to the ovule, which is deeply embedded in the pistil base^2^. To ensure successful fertilization, angiosperms have evolved multiple mechanisms for pollen–pistil interactions involving ovular chemotropism^3^. Chemotropism is the directed growth of cells, tissues, or organisms in response to external cues, such as the axon guidance of olfactory sensory neurons expressing individual odorant receptors^4, 5^. Angiosperms employ a chemotropic one-to-one guidance system known as “pollen tube guidance.” *Arabidopsis thaliana* flowers possess single ovaries comprising dozens of linearly aligned ovules (Fig. S1A). After the pollen tube enters the transmitting tract (TT), it reaches the ovule via three steps: emergence from the TT, growth on the funiculus (funicular guidance), and penetration of the micropyle (micropylar guidance). Genetic studies have demonstrated that ovules regulate micropylar guidance via the secretion of small defensin-like peptides (LUREs), which interact with receptor-like kinases on the pollen tube^6–9^.

Egg fertilization by more than one sperm (polyspermy) is restricted to maximize reproductive success^10^. The prevention of multiple pollen tubes penetrating a single ovule is termed “polytubey blocking”^11^. Since ovule-derived chemoattractants can affect all pollen tubes, it is thought that an ovule can potentially attract multiple pollen tubes. However, polytubey is rare in *A. thaliana* (∼1%)^12, 13^, despite the presence of ∼60 ovules and >60 pollen tubes^14^. Recently, multistep polytubey blocks have been reported. A FERONIA-dependent block located in the septum affects pollen tube emergence^15^. When a pollen tube arrives at the micropyle, FERONIA in the gametophytic synergid cells triggers the accumulation of nitric oxide, which inactivates LUREs^16^. Due to the lack of LURE-type attractant peptides following gamete fusion, no further pollen tubes are attracted^17–19^. However, because the real-time behavior of pollen tube attraction has not been observed, the full context of the blocking system remains unclear. Here, we developed an imaging method using two-photon excitation microscopy with the aim of investigating pollen tube dynamics in living ovaries of *A. thaliana*.

## RESULTS

### Two-photon Imaging Revealed Pollen Tube Dynamics in a Living Ovary

In the wild-type (WT) *Col-0* ecotype of *A. thaliana*, the average number of pollen tubes growing in the TT at 18 hours after pollination (HAP) was 70 ± 3.6 following maximum pollination (n = 3 pistils; Fig. S1B−E1). The behavior of each pollen tube in the fixed tissue was difficult to analyze (Fig. S1B). We developed a “single-locule method” for deep imaging of pollen tube dynamics inside a live *Arabidopsis* ovary expressing fluorescent reporter (Fig. S21). In the WT, pollen tubes that entered the ovary reached its base at 16 HAP (Movie S1A, Fig. 1A−C). The pollen tube growth rate was not constant, and gradually reduced to 1.3 ± 0.1 μm/min (n = 7 pollen tubes) between 4 and 16 HAP (Fig. S3A). Some pollen tubes abruptly separated from the pollen tube bundle to pass the septum epidermal (SE) layer, adjusting their growth towards the locule (Fig. 1C). Such directional changes were assumed to indicate pollen tube emergence, because the fluorescent signals of these tubes showed that they elongated along the funiculus after passing through the SE (Fig. S2C), being attracted to the fluorescent-labeled synergid cells (Fig. 1F–G, Movie S1B). To our knowledge, this is the first report of such dynamic pollen tube growth in the ovary. Our single-locule method enabled a real-time analysis of pollen tube guidance until it reached the micropyle.

**Fig. 1.**
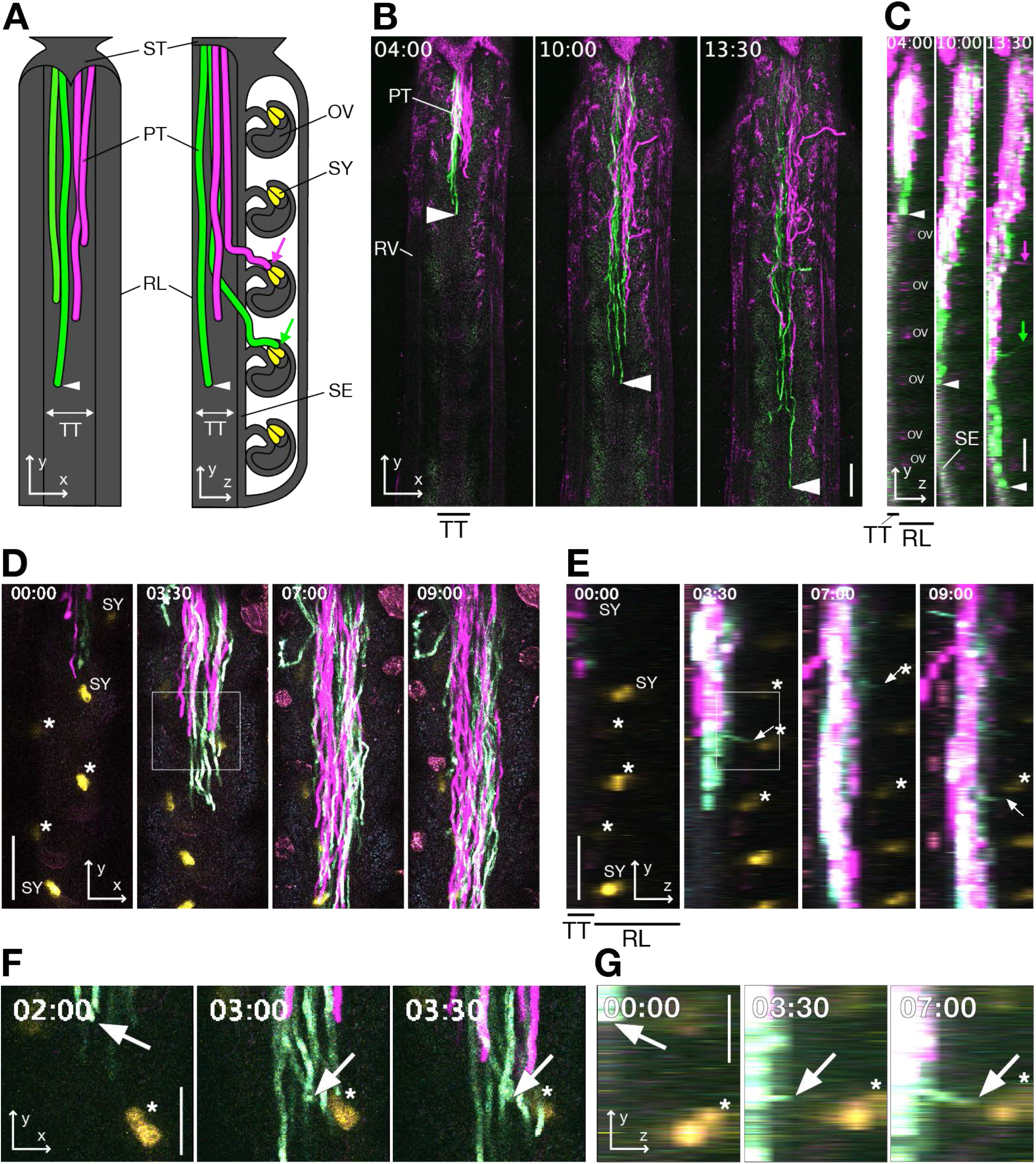
Pollen tube dynamics in the *Arabidopsis* living pistil. (A) Schematic representations of the *xy-* (left) and *yz-* (right) images of pollen tube growth in the ovary under the single-locule method. (B and C) WT pistil pollinated with mixture of pollen expressing mTFP1 (green) and TagRFP (magenta). The *xy-* (B) and *yz-* (C) maximum projection images by 10-μm steps including TT were shown. Magenta and green arrows indicate pollen tubes guided toward the ovule after emergence from the TT. (D and E) The pistil having ovules with synergid cells labeled by GFP (yellow) was pollinated with the pollen expressing mTFP1 (cyan) and TagRFP (magenta). The *xy-* (D) and *yz-* (E) maximum projection images by 7 z-stack images with 15-μm steps were shown. (F and G) Magnified images of the funicular guidance highlighted by a white box in (D) and (E). Arrowheads indicate the tip of most growing pollen tubes. Arrows indicate pollen tubes guided toward the ovule after emergence from the TT. Asterisks indicate synergid cells of the target ovules. The numbers stamped in each frame indicate time (hh:mm) from the start of the observation. ST, style; PT, pollen tube; RL, remaining locule; TT, transmitting tract; OV, ovule; SY, synergid cell; SE, septum epidermis. Scale bars, 100 μm (B–E) and 50 μm (F and G). See also Movie S1.

### Ovule Targeting Depends on the Number and Distribution of Pollen Tubes

In *A. thaliana*, it remained unclear which pollen tube would selectively be attracted, and which ovule would preferentially attract a pollen tube^20, 21^. We investigated the parameters determining the priority of pollen tube attraction. Under maximum pollination, pollen tubes were distributed throughout the entire TT (Fig. S1B–E); however, following limited pollination (Fig. 2A–C), available space allowed the tubes to grow freely anywhere in the TT (Fig. 2G–J). Under maximum pollination, the median distribution of the targeted ovules was 22.0% (205 ovules in 12 pistils, Fig. 2F), which were distributed towards the apex. Utilizing limited pollination, a single pollen grain was applied to a stigma (Fig. 2A and 2B). The single seed distribution in the silique indicated that the germinated single pollen tube reached it (Fig. 2C). The median single seed distribution from the top of the silique was 49% in for central pollination (n = 52 siliques, Fig. 2D) and 46% for side pollination (n = 44 siliques, Fig. 2E). The apical 10% of ovules were not targeted in either condition (Fig. 2D and 2E). Therefore, preferential ovule targeting depended on the number of germinated pollen grains. Moreover, the results of this experiment suggested that a pollen tube can pass apical ovules, implying that an ovular attraction signal may only be effective locally. Pollen tube behavior and growth path was also investigated using an SE reporter to determine which pollen tube emerged from the TT^22^ (Fig. S2B and S2C). Under limited pollination, some pollen tubes gradually moved toward the SE and emerged from the TT, whereas other pollen tubes grew down the TT (Fig. 2G and 2H). When two pollen tubes elongated in parallel, the one closer to the SE was more likely to emerge (Fig. 2I and 2 J). Therefore, we concluded that pollen tubes were attracted from the top down in maximum pollination but not in limited pollination, for two reasons: because pollen tubes adjacent to the SE were selected for emergence in the TT, and because in maximum pollination, pollen tubes were more likely to grow adjacent to the SE.

**Fig. 2.**
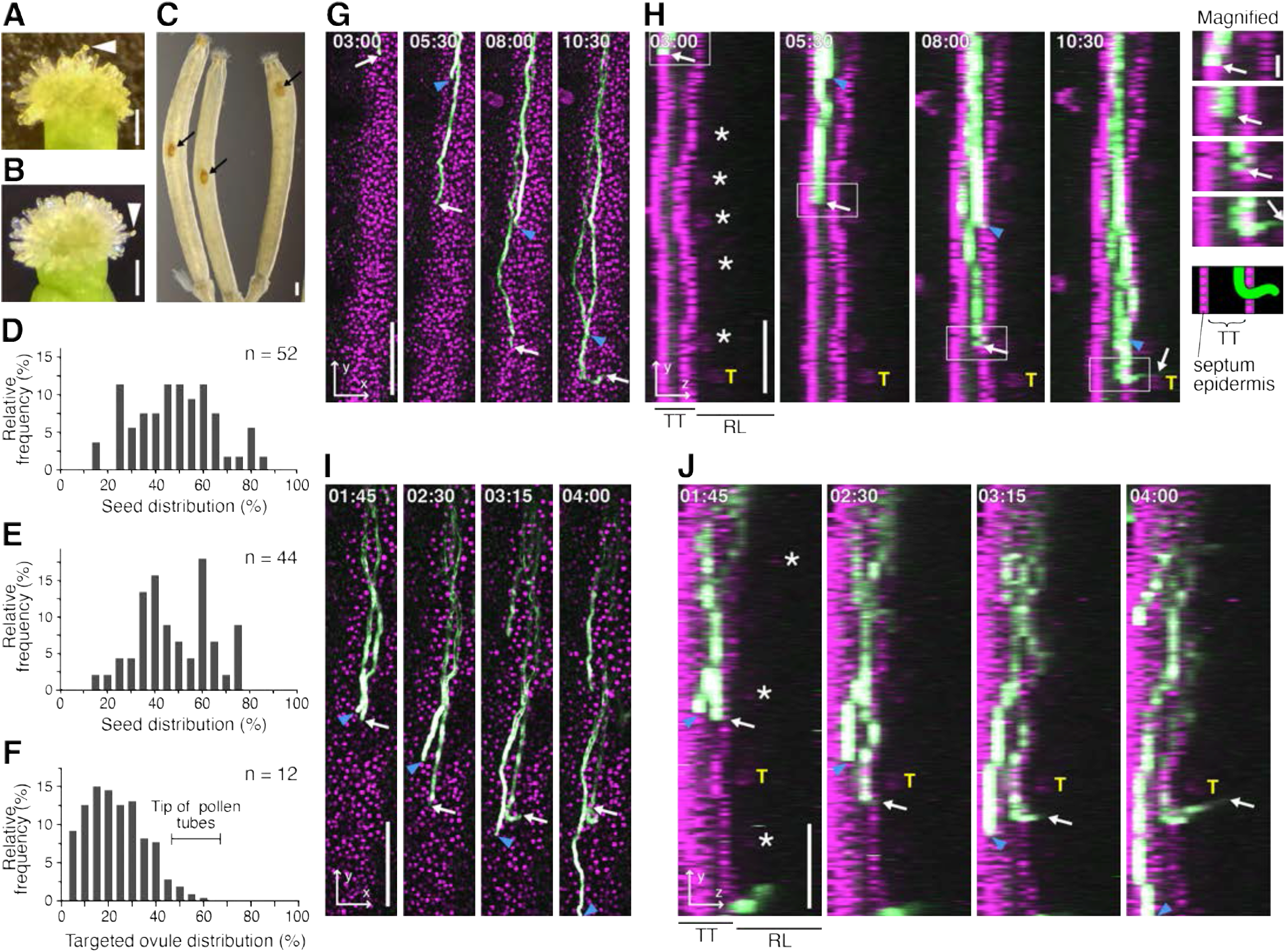
Emergence and ovule targeting of the pollen tube from the transmitting tract under limited pollination. (A and B) Spatially regulated limited pollination. Pollinated single pollen grains are shown as arrowheads at the center (A) or side (B) of the stigma. (C) Transparent siliques with one seed from each pollination of single pollen. Seeds were indicated by arrows. (D and E) Single seed-set distribution of the 52 pistils with center pollination (D) and 44 pistils with side pollination (E) under the single pollen pollination. Bar charts represent the frequency of seeds present in each of the ten percentiles of locule length, from the most apical (0%–10%) to the most basal (90%–100%) of each locule. (F) The distributions of the 205 targeted ovules in the maximum pollinated pistils, when the tip of pollen tube reached at the middle part at 4 h after pollination (n = 12 pistils, 56.5 ± 7.1%). (G–J) Emergence of the pollen tube from the transmitting tract. Pollen tubes and nuclei of the septum epidermis labeled with each fluorescent protein are shown in green and magenta, respectively. See also Movie S2A and S2B. The *xy-* (G and I) and *yz-* (H and J) projection images by 10-μm steps with 18 planes. Magnified images with schematic representation of the emergence of pollen tube from the transmitting tract highlighted by a white box in (H). (I and J) Pollen tube at the closest to the septum epidermis emerged from the transmitting tract. Arrows and arrowheads indicate emerged and non-emerged pollen tubes. Asterisks show the autofluorescence of ovule. Time stamps are the same as in Figure 1. T, targeted ovule; TT, transmitting tract; RL, remaining locule. Scale bars, 100 μm, and 20 μm in the magnified image of (H).

### Ovular Sporophytic Tissue Controls the Behavior of Pollen Tubes Inside the TT and Their Emergence From It

To assess properties of the emergence signal(s) in the TT, we investigated the temporal regulation of pollen tubes that had emerged. Two-photon live imaging showed that the growth rate of emerged pollen tubes gradually decreased to 0.5 μm/min approximately 2 h before their emergence (Fig. 3A); non-emerged tubes maintained a growth rate of approximately 1.4 ± 0.3 μm/min (n = 69 pollen tubes in 10 pistils). The *xy*- and *yz*- projection images revealed that the pollen tube growth rate decreased following its attachment to the SE inside the TT (Fig. 3B−C and Fig. S3B). The median time from pollen tube SE attachment until emergence was 120 min (n = 11; range 45–285 min), consistent with the decreased growth rate (Fig. 3A). This suggested that SE attachment inside the TT triggers a reduction in the pollen tube growth rate, persisting throughout its emergence.

**Fig. 3.**
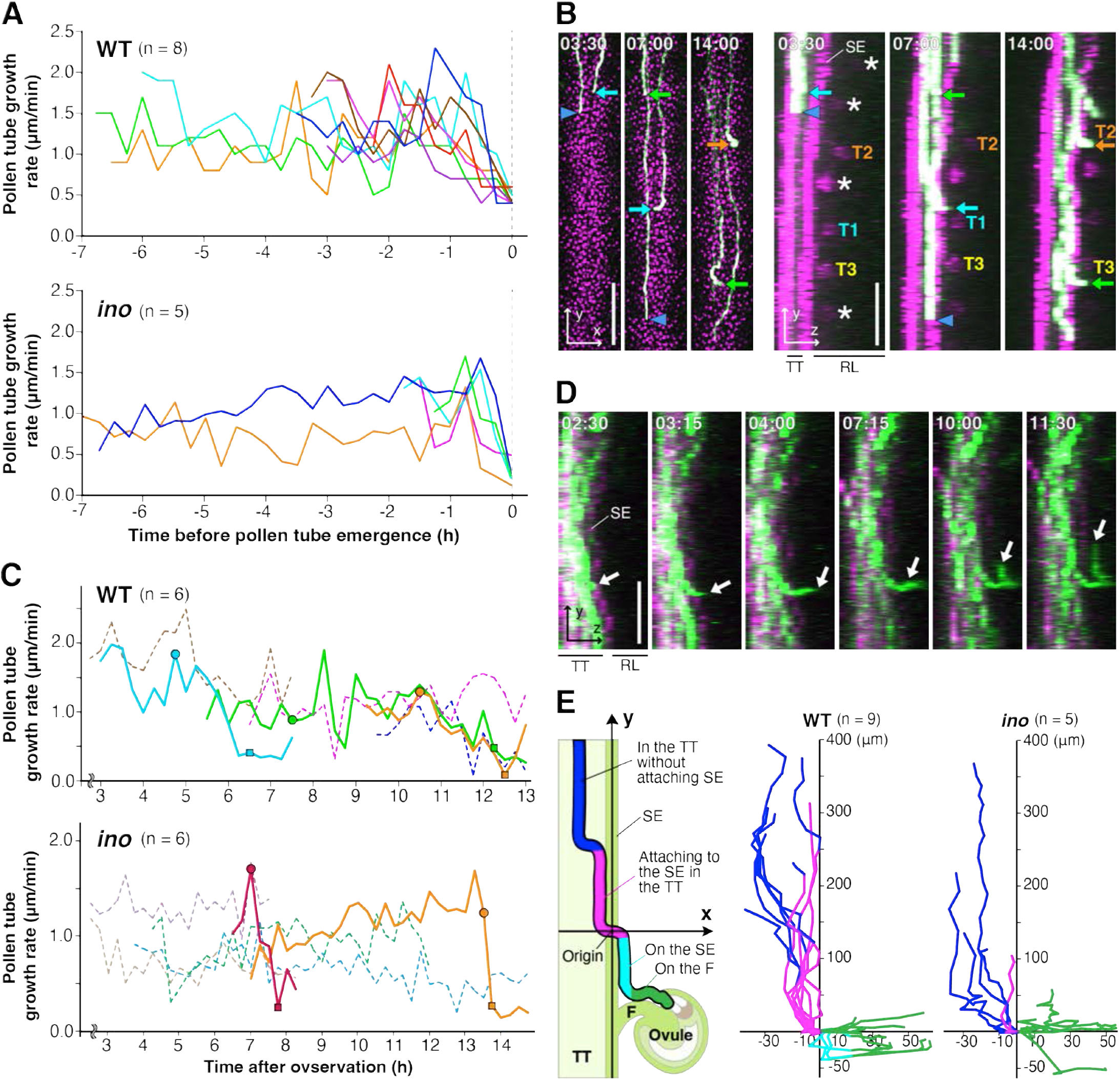
Features of emerged pollen tube from the transmitting tract in the WT and *ino* mutant ovaries. (A) Pollen tube growth rate in the transmitting tract of 8 and 5 pollen tubes that emerged from the transmitting tract in the WT and *ino* pistils, respectively. Time of pollen tube emergence was defined as origin. A value of x smaller than zero indicates pollen tube within the transmitting tract. (B) Emergence of the three pollen tubes from the transmitting tract in the WT ovary. Asterisks show ovule autofluorescence. Arrows and arrowheads indicate emerged and non-emerged pollen tubes, respectively. Each targeted ovule is shown as T1–T3. See also Movie S2C. (C) Pollen tube growth rate of emerged and non-emerged pollen tubes in a single pistil is shown as solid and dashed lines respectively. Filled circles show the moment of pollen tube attachment to the septum epidermis inside the transmitting tract. Filled squares show the moment of pollen tube emergence. (D) Pollen tube emerging from the transmitting tract of the *ino* pistil. White arrows indicate the tip of emerged pollen tube. (E) Tracking of the tip of 9 and 5 emerged pollen tubes in the WT and *ino* pistils from the *yz*-projection images. The emerged pollen tubes ware colored by its *yz*-positions as follows: growing in the transmitting tract (blue), attaching to the septum epidermis in the transmitting tract (magenta), growing on the septum epidermis after the emergence (cyan), and growing on the funiculus (green). Time stamps are the same as in Figure 1. SE, septum epidermis; F, funiculus; OV; ovule; TT, transmitting tract; RL, remaining locule. Scale bars, 100 μm.

Ovular cells are indispensable for ovule targeting^23, 24^. To investigate the effect of ovule maturity on ovule targeting, ovule development was investigated in ClearSee-cleared pistils^25, 26^. Ovule maturation was indicated with the appearance of a fluorescent signal of the FGR8.0 reporter, which combines ovular cell fluorescent markers^17^. Maturation commenced in the middle to lower portion of the ovary at stage 11, then extended to both the upper and lower extremities until stage 14 (Fig. S4A). Even on the day of flowering, at stage 14, some ovules in the topmost ovary remained immature (arrows in Fig. S4A). This result correlated well with the ovule targeting preference under limited pollination conditions (Fig. 2), implying that ovule maturation contributes to ovule targeting.

To investigate how ovular cells are involved in targeting, ovular mutants pollinated with WT pollen were examined. In the WT, pollen tubes emerging from the TT adhered to the septum surface^27^, then elongated along the funiculus toward the ovule (Fig. 4A). The *ant* ovary exhibits a complete loss of megasporogenesis, resulting in severe ovular developmental defects^28^. The *ino* mutant lacks outer ovule integuments^29^. In both mutant ovaries, there was a reduced number of emerged pollen tubes and ovules showing funicular guidance (Fig. 4B). A disorder of pollen tube adhesion to the septum and funicular surfaces was also observed (Fig. 4A and 4 B). The *dif1* mutant lacks an embryo sac due to meiotic chromosome mis-segregation^30^. Although micropylar guidance was completely impaired in the *dif1* ovary, pollen tubes emerged from the TT and adhered to the female tissue surface, whereafter many of them lost their way to the micropyle and continued their disordered growth in the ovary locule (Fig. 4A and 4 B). The gametophytic mutant *myb98* possesses defective synergid cell differentiation in the embryo sac^31^. The number of pollen tubes emerging in such ovaries was reduced, and they showed adhesion but sometimes non-attachment to the female tissue surfaces (Fig. 4A and 4 B). Our results suggest that the sporophytic ovular outer integument enhances pollen tube emergence and adhesion to the maternal tissue surface. To confirm this, pollen tube emergence in the *ino* ovary was assessed through two-photon live imaging. In the WT, pollen tubes gradually changed their growth direction inside the TT, then elongated along the SE for approximately 50–300 μm before emergence (Fig. 3A). In contrast, pollen tubes in the *ino* TT did not grow along the SE (Fig. 3E) and emerged rapidly without decreasing their growth rate (Fig. 3A and 3C), with disordered growth after emergence from the TT (Fig. 3D). The *Arabidopsis* TT comprises longitudinally elongated cells and a large amount of extracellular matrix components^20^, but there were no obvious morphological differences between the TTs of WT and *ino* (Fig. S4B). These results suggest that a signal(s) derived from the ovular outer integument affects pollen tube behavior, even inside the TT, which most likely facilitates pollen tube emergence.

**Fig. 4.**
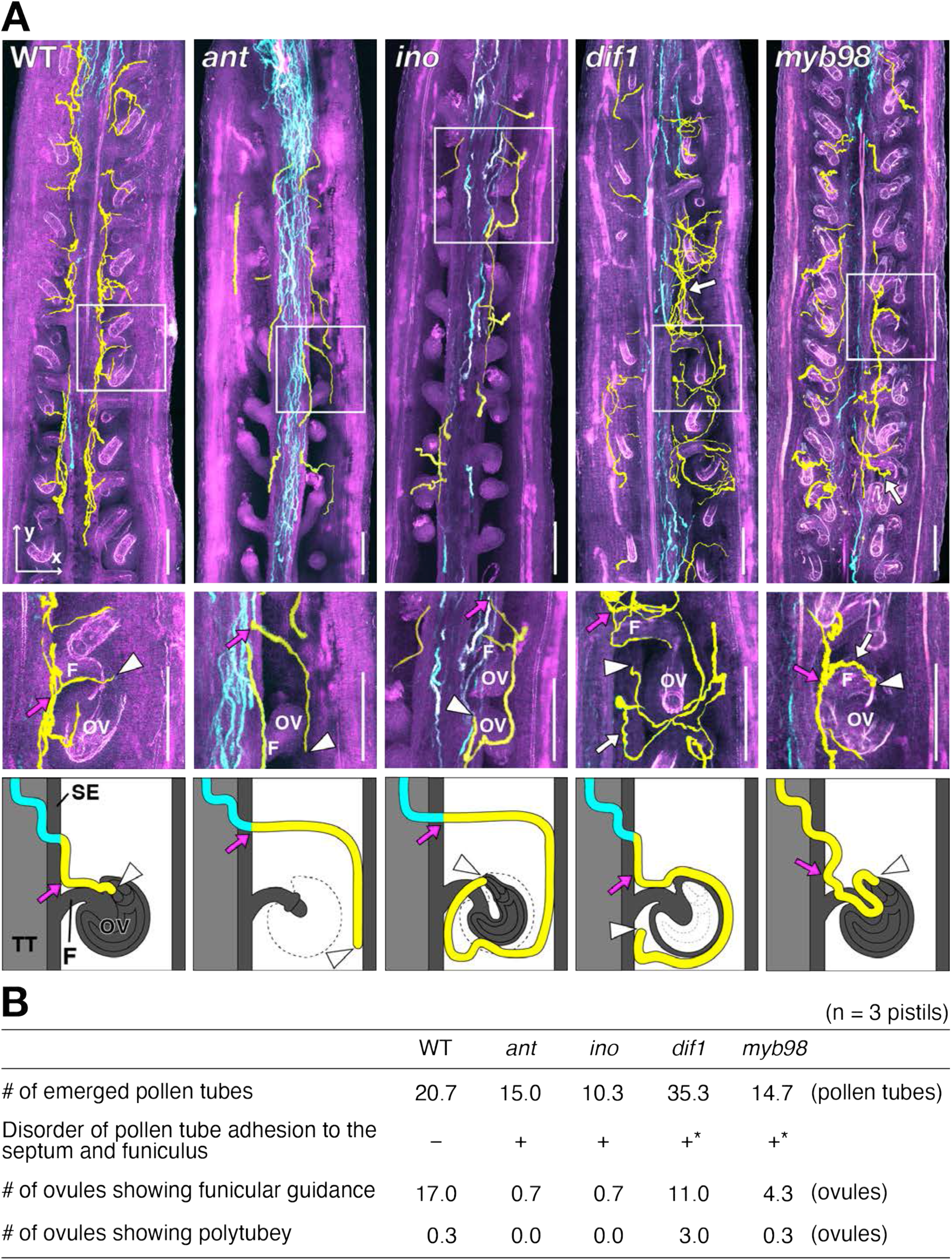
Pollen tube emergence and funicular guidance in the four mutant pistils showing impaired ovular tissues. (A) Observation of pollen tube behavior in the ovary of ClearSee treated maximum pollinated pistils. The pistils pollinated with pollen labeled by mTFP1 was collected 12– 18 hours after pollination. The *xy-*projection images by 8-μm steps were shown. Pollen tubes expressing mTFP1 and pistil autofluorescence were shown in cyan and magenta. The pollen tubes that emerged from the transmitting tract into the locule are colored with yellow and overlaid. Magnification of emerged pollen tube highlighted by a white box and its schematic representation are also shown at the bottom. Arrows and arrowheads indicate the point of pollen tube leaving from the septum and the tip of pollen tube, respectively. F, funiculus; OV, ovule; SE, septum epidermis; TT, transmitting tract. Scale bars, 100 μm. (B) Features of emerged pollen tubes in the WT and four mutant ovaries. Pistils pollinated by pollen labeled with mTFP1 and Venus were collected at 6 h after pollination and cleared by ClearSee. Three pistils per each mutant were averaged. Asterisks show pollen tube adhesion but sometimes non- attachments to the female tissue surfaces (white arrows). Ovule having pollen tubes on the funiculus was defined as funicular guidance. Ovule having multiple pollen tubes on the funiculus was counted as polytubey.

### The Regulation of Pollen Tube Emergence Facilitates Polytubey Blocking

In *feronia* (*fer*) and *lorelei* (*lre*) mutant pistils, pollen tube reception by synergid cells is impaired^12, 32^, resulting in polytubey (Fig. 5A). The homozygous *gcs1/hap2* male mutant impairs gamete fusion, which results in high-frequency polytubey^33^ due to the fertilization recovery system^34^. Multiple pollen tubes overlapped and were attracted to the same route as the funiculus in *gcs1*, whereas elongated tubes did not overlap on the funiculus in most *fer* and *lre*, along with a few WT ovules (Fig. 5A), as was previously reported in *gcs1*^35^ and *magatama* mutants^36^. This suggests a restart of pollen tube guidance for fertilization recovery (*gcs1*/*hap2*), and that polytubey before reception of the first tube (*fer*, *lre*, and WT) involves different mechanisms in the attraction and/or repulsion signal(s) on the funiculus^36^.

**Fig. 5.**
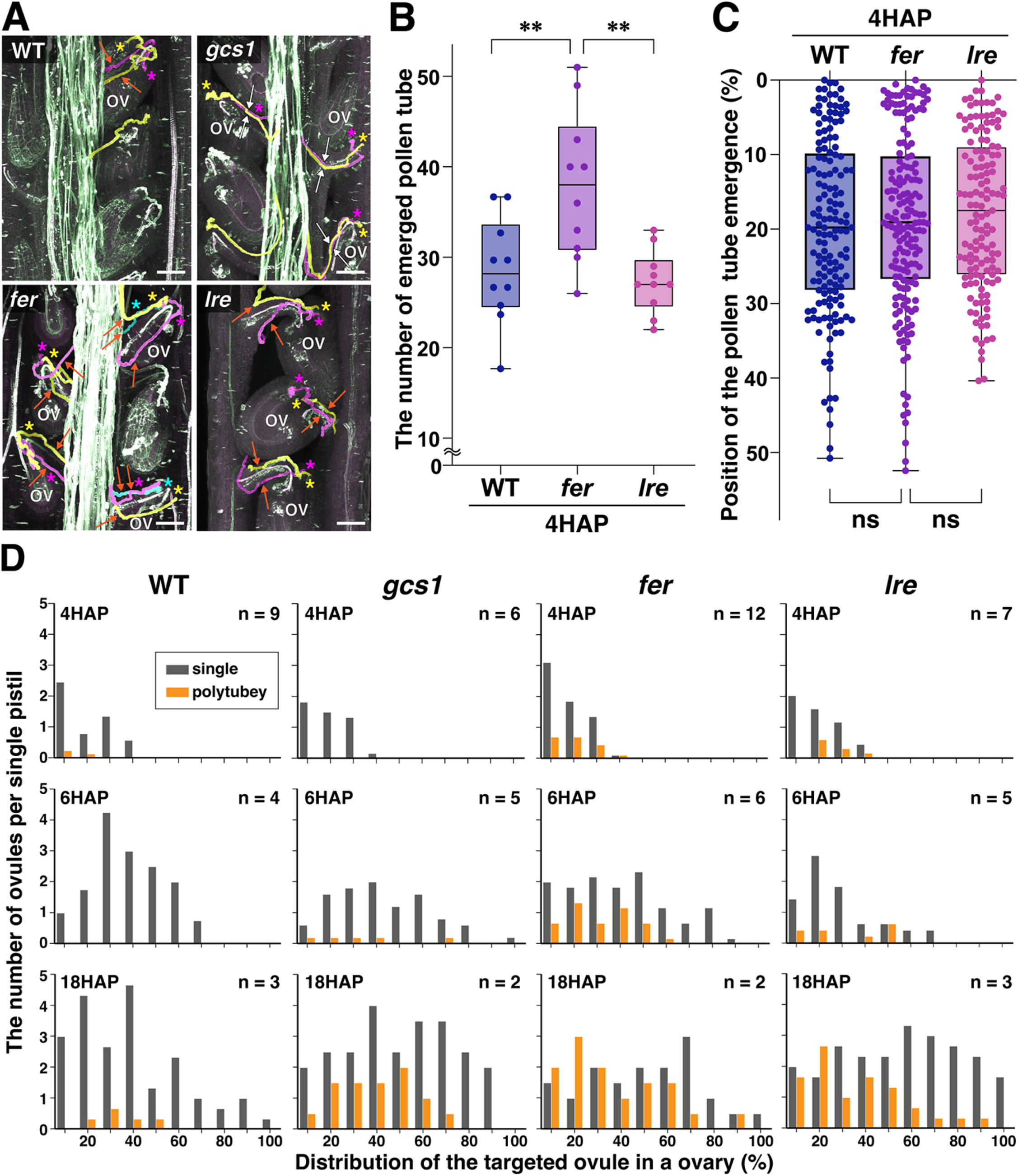
Spatiotemporal analysis of pollen tube emergence and funicular guidance in the cross showing polytubey by using *fer*, *lre*, and *gcs1* mutants. (A) Funicular guidance in the WT and mutant ovaries. Pollen tubes in the pistil were stained with aniline blue in the NaOH-clearing solution. Projection images were generated from 2-μm steps. Multiple pollen tubes adhering on the same funiculus were colored overlay with yellow, magenta, and cyan with same-colored asterisks. Multiple pollen tubes that overlapped at the same route on the funiculus are shown as white arrows, whereas those that did not overlap on the funiculus are shown as orange arrows. OV, ovule showing polytubey. (B) The number of emerged pollen tube in the pistils of WT, *fer*, and *lre* at 4 HAP by 10 independent experiments (n = 10 pistils). (C) The position of the pollen tube emergence on the septum epidermis at 4 HAP in the pistils of WT, *fer*, and *lre* by sum of each 5 pistil. (B and C) Box plots represent the median with 25th and 75th percentiles with minimum and maximum whiskers. (D) The number of ovules showing single pollen tube attraction (gray) and polytubey (magenta) per ovary at 4, 6, and 18 hours after pollination (HAP). Bar charts represent the number of ovules present in each ten percentile of ovary length from the most apical (0%–10%) to the most basal (90%–100%) of each pistil. Statistical significance was determined using one-way ANOVA followed by Tukey’s multiple comparisons tests. ∗∗p < 0.01; ns, not significant. Scale bars, 50 μm.\

FER is widely expressed in female tissues and is involved in polytubey blocking by both the sporophytic tissues and synergid cells^15, 16^. In contrast, LRE shows specific expression in synergid cells^37^. To investigate polytubey blocking during pollen tube emergence and funicular guidance, pollen tube emergence was compared among WT, *fer*, and *lre* pistils at 4 HAP, when fertilization is incomplete in most ovules. The number of emerged pollen tubes was significantly increased in the *fer* pistil, even at 4 HAP, but not in the *lre* pistil (Fig. 5B). However, the position of pollen tube emergence on the septum surface did not differ significantly among groups (Fig. 5C). We deduced that polytubey prevention via regulation of the number of pollen tubes emerging was already in effect at 4 HAP in the WT but was impaired in *fer*.

The distribution of ovules showing polytubey in mutants was investigated in NaOH-cleared pistils. Polytubey was scarce among WT ovules but prevalent in *fer* and *lre*, even at 4 HAP (Fig. 5D). Polytubey was more severe in *fer* than in *lre*, suggesting that regulation at the emergence step contributes significantly to polytubey block on the funiculus. In contrast with WT and *gcs1* ovules, those of *fer* and *lre* showed polytubey at 18 HAP in the upper part of the ovary. Overall, polytubey phenotypes on the funiculus were similar in *fer* and *lre* mutants (Fig. 5A and 5D), but the number of emerged pollen tubes differed between them (Fig. 5B). Therefore, polytubey block may be regulated by FER during both pollen tube emergence and funicular guidance, but only during funicular guidance by LRE.

### Entry of the Second Pollen Tube onto the Funiculus Is Rapidly Blocked by Gametophytic Signals

The spatiotemporal behavior of pollen tubes during polytubey was observed under the single-locule method. Across 110 WT pistils, 448 ovules displayed pollen tube attraction of which 18 possessed polytubey, representing a rate of 4.0% (Fig. 6A and Movie S3A). Polytubey was more frequently observed in *lre* (19 pistils: 71 ovules displaying attraction, 35 with polytubey; a rate of 49%) and *fer* (27 pistils: 111 ovules exhibiting attraction, 37 with polytubey; a rate of 33%) than in the WT (Fig. 6B and 6C). In both mutants, several pollen tubes returned from the funiculus to the septum (i.e., repulsion), but none did in the WT (arrows in Fig. 6B and 6C, Movie S3B and S3C). A repulsive signal may therefore be inhibiting the funicular guidance of excess pollen tubes, which is active in both *fer* and *lre* ovaries.

**Fig. 6.**
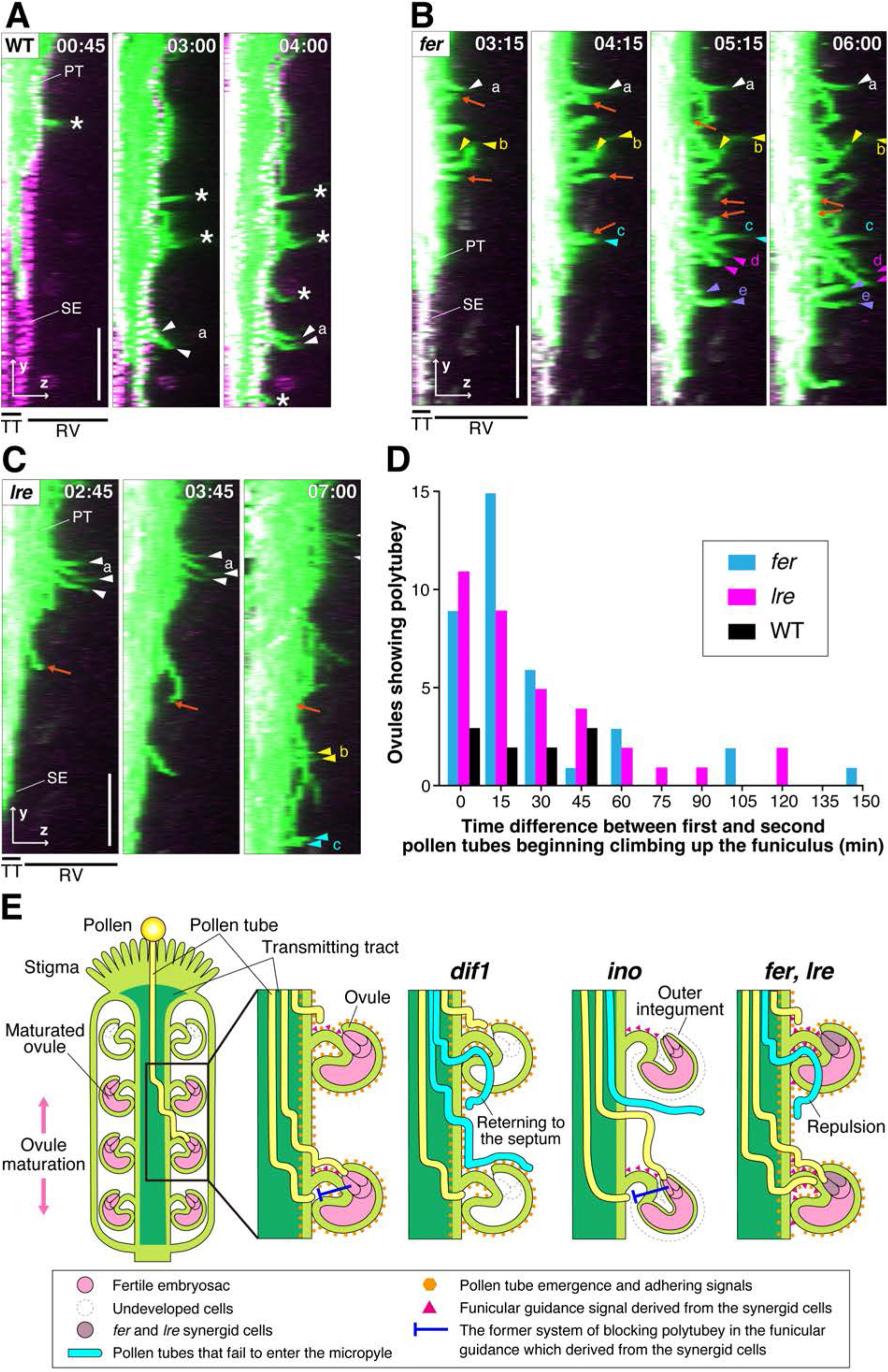
Live imaging of polytubey in the WT and mutant ovaries and model of one-to-one pollen tube guidance. (A–C) Polytubey in WT, *fer*, and *lre* pistils pollinated with mApple-expressing pollen. The *yz*-maximum projection images in 7- or 8-μm steps, including the transmitting tract, are shown. The pollen tube and pistil autofluorescence are shown in green and magenta, respectively. Asterisks indicate single-pollen tube guidance. Pollen tubes on the same funiculus are shown as same-coloured arrowheads. Small letters indicate ovules with polytubules. Arrows indicate pollen tubes that leave the septum but return to the septum surface. The numbers stamped in each frame indicate the time (h:mm) from the start of observation. Scale bars, 100 μm. See also Movie S3. (A) WT pistil with nuclei of the septum epidermis labelled by NLS-YFP is shown in magenta. (B) The pistil of *fer* mutant. (C) The pistils of *lre* mutants. (D) Time difference between the first and second pollen tubes beginning climbing up the funiculus in the case of polytubey. The number of ovules showing polytubey in the WT (n = 10 ovules), *fer* (n = 37 ovules), and *lre* (n = 35 ovules) pistils is summarized. The funicular guidance time for each first pollen tube was set at 0 min. (E) Model of one-to-one pollen tube guidance spatiotemporally regulated by female sporophytic and gametophytic cells. When the number of pollen tubes is limited, they are preferentially attracted to the ovule located in the lower center of the ovary, depending on the ovule maturation. The pollen tube emergence signal was derived from ovular sporophytic cells. This signal affects the attachment of pollen tubes to the funicular and ovular surfaces and the septum epidermis inside the transmitting tract, which causes a decrease in growth rate and pollen tube emergence into a locule. The emerged pollen tube elongates with attachment to the septum surface and receives funicular guidance signals in a spatially restricted region of the funiculus. The signals derived from the sporophytic and gametophytic cells prevent multiple pollen tube attractions, blocking polytubey in funicular guidance. In the *dif1* ovary, the pollen tube emergence signal from the ovular outer integument was normal, but gametophytic cell-dependent blocking polytubey was impaired. In the *ino* ovary, which lacks pollen tube emergence and adhesion signals, a few lucky floating pollen tubes arrive at the micropyle because the signals derived from gametophytic cells work normally. Multiple pollen tubes are attracted along different paths on the funiculus in the *fer* and *lre* ovaries because the gametophytic-dependent attractive funicular guidance signal is increased and/or expanded, or the hypothetical repulsive signal decreases and/or narrows. Pollen tubes that failed to enter the micropyle returned to the septum and then attempted to head for another ovule in the *dif1*, *fer*, and *lre* ovaries. The blocking system for preventing polytubey during funicular guidance is regulated by both sporophytic and gametophytic cells. Signals derived from gametophytic cells were impaired in *dif1*, *fer*, and *lre* mutants, which causes polytubey.

Next, the timing of funicular guidance initiation between the first and second pollen tubes was analyzed. All 10 polytubey events observed in the WT (in which the second pollen tube was attracted without repulsion) were observed within 45 min (Fig. 6D). Similarly, 31 of 37 (84%) and 29 of 35 (83%) polytubey events were observed within 45 min for *lre* and *fer*, respectively. These results suggest that the polytubey block is not strictly prohibitive for 45 min after entry of the first pollen tube onto the funiculus, even in the WT. The frequent polytubey within 45 min in both *fer* and *lre* suggests that the distantly locating synergid cell is involved in rapid but non-strict blockage of entry onto the funiculus. Heterozygous *fer* and *lre* mutants showed frequent and comparable polytubey on the funiculus at 6 HAP, suggesting that it is not the sporophytic but the gametophytic signal which is needed for blockage of second pollen tube entry onto the funiculus (Fig. S4D). After 45 min, some instances of attraction of the second pollen tubes were observed in the *fer* and *lre* ovaries (Fig. 6D). Overall, the polytubey block was stricter 45 min post entry of the first pollen tube, and the defect in the FER and LRE signaling pathway of the female gametophyte (possibly in the synergid cell) may allow polytubey during this period.

## DISCUSSION

*A. thaliana* pistils contained ∼60 ovules, but less than two times the number of pollen tubes grew in the pistil (Fig. S1E). Thus, one-to-one pollen tube guidance is critical for effective reproduction. Our live imaging suggested that the first step in this process was selection within the TT, which depended on pollen tube distribution. This suggests that selection is stochastic rather than dependent on pollen tube capacitation^38^, and that the emergence signal(s) is not uniform throughout the TT but is more effective in the vicinity of SE. Pollen tube emergence was normal in *dif1* but decreased in *ant* and *ino* mutants. Ovular outer integument-dependent signals may enhance pollen tube emergence and regulate adherence to the maternal tissue surfaces. These signals are likely to be localized to the surfaces, but they may also reach inside the TT (Fig. 6E). The more mature the ovule, the deeper into the TT the signal(s) for emergence might reach.

Both gametophytic signals^39^ and regulatory sporophytic molecules have been associated with pollen tube guidance^40^, for example exogenous gamma-aminobutyric acid^41, 42^, arabinogalactan proteins^43^, ovular methyl-glucuronosyl arabinogalactan^44^, the ethylene precursor 1-aminocyclopropane-1-carboxylic acid^45^, and ovary-expressed bHLH transcription factor^46^. However, the nature of directional cues from ovular sporophytic tissue remains unclear. Molecules directly involved in the adhesion of pollen tubes to female tissues are undescribed, although stylar cysteine-rich adhesin (SCA) is known to enhance the attraction activity of chemocyanin and adhesion to the lily pistil^47, 48^. Our findings should accelerate the identification of ovular outer integument-derived signaling molecules involved in pollen tube adhesion and attraction.

Polytubey blocking is achieved in multiple steps, and mutants lacking them have been studied, including *myb98*, *fer*, *lre*, *maa1*, *maa3*, and *gcs1*^31, 32, 34, 36, 49^. *FER*, *ANJ*, and *HERK1* receptors on the septum interact with pollen tube-produced RALF peptide ligands, which are involved in polytubey blocking^15^. We observed a blocking defect in the *fer* ovary, which exhibited a higher number of pollen tubes, although the location of their emergence remained unchanged. The polytubey block is achieved by preventing pollen tube entrance into the micropyle through multiple steps, including dispersion, modification, and degradation of pollen tube-targeting chemoattractants^16–19^. Through our novel imaging approaches, we discovered that regulation also occurs earlier, during entry on the funiculus (Fig. 6E). A block was rapidly established following entry of the first pollen tube, dependent on gametophytic FER and LRE. In the first 45 min, this block was less restrictive, even in the WT. This agrees with genetic experiments showing that polyspermy in *Arabidopsis* can be explained by fertilization via more than one pollen tube^50–53^. Polyspermy blocking exists in plant gametes, although the block is more strict and rapid in the egg than the central cell^54, 55^. On the egg cell, an almost simultaneous pollen tube discharge is required for polyspermy^51^, but its mechanistic basis is unclear under one-to-one pollen tube guidance^11^. Our live imaging suggests that two pollen tubes simultaneously approaching the same ovule within 45 min may induce polyspermy in *Arabidopsis*.

Molecular properties of the polytubey block upon pollen tube entry onto the funiculus remain elusive. In the synergid cell, FER-dependent signaling mediated by de-esterified pectin induces NO production, which inactivates LUREs^16^. Our results imply long-distance communication between the first pollen tube and the synergid cell, because blocking commenced before the first tube arrived at the micropyle (Fig. 6D). Molecular diffusion before and after 45 min requires further research. Our live imaging also revealed the novel repulsion process of excess pollen tubes on the funiculus, the mechanism of which remains largely unknown^23^. This repulsion mechanism is likely to be FER- and LRE-independent because it occurred in *fer* and *lre* more frequently.

In conclusion, our two-photon imaging revealed novel dynamics and spatiotemporal signaling in one-to-one pollen tube guidance. This should accelerate the identification of signaling molecules driving this process in angiosperms, which exhibits a unique navigation system related to sexual reproduction.

---

**Fig. S1.**
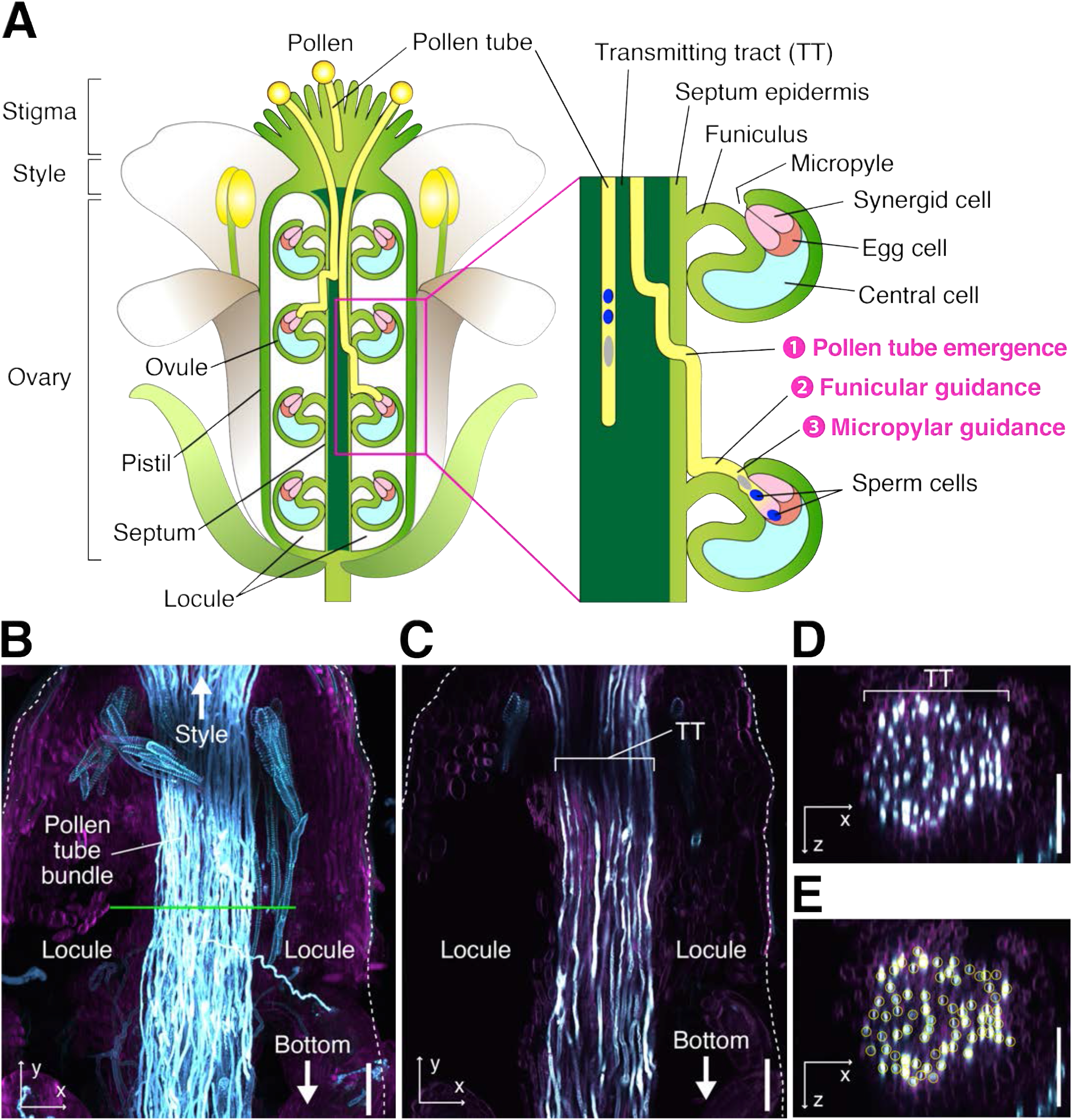

**Fig. S2.**
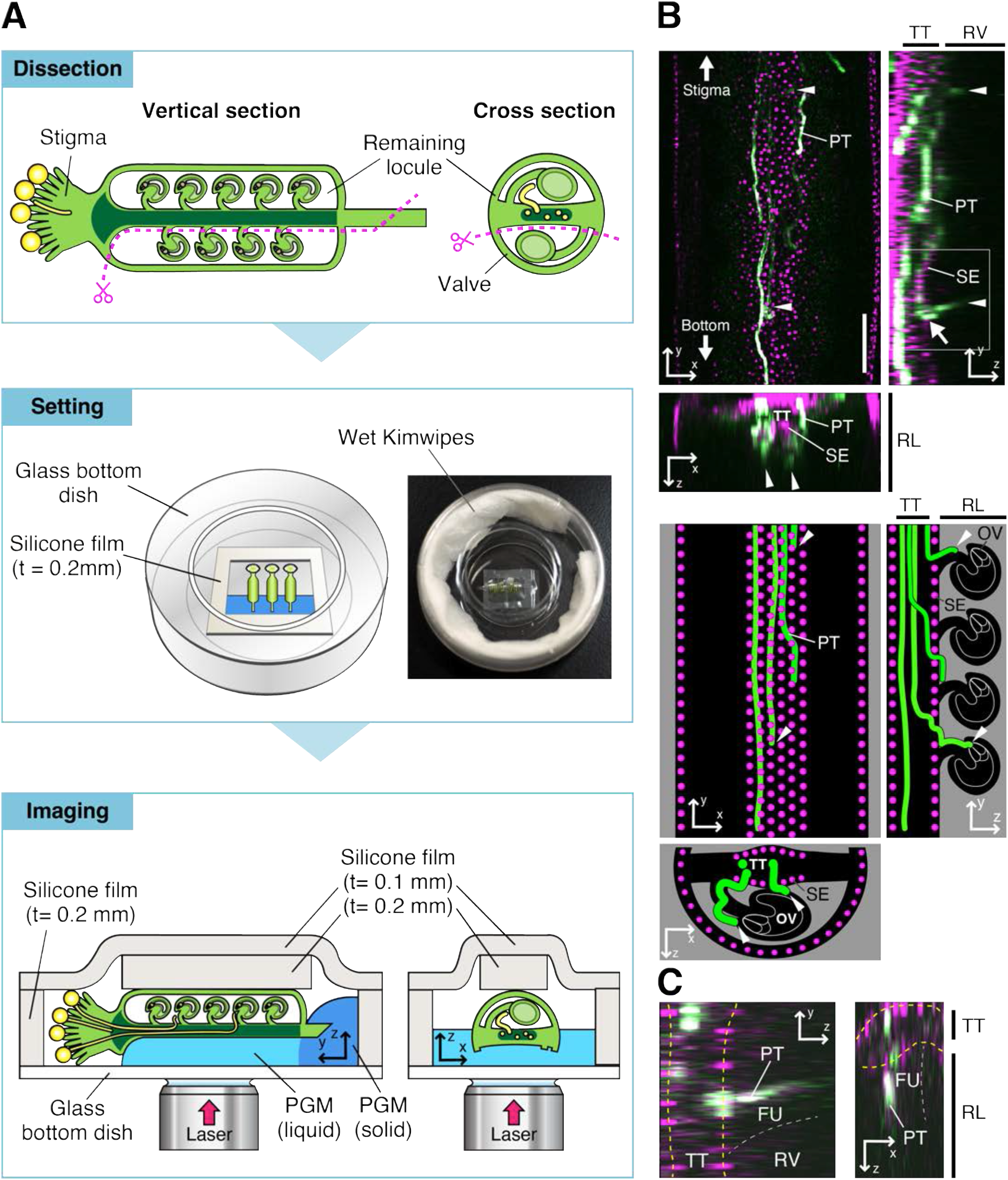

**Fig. S3.**
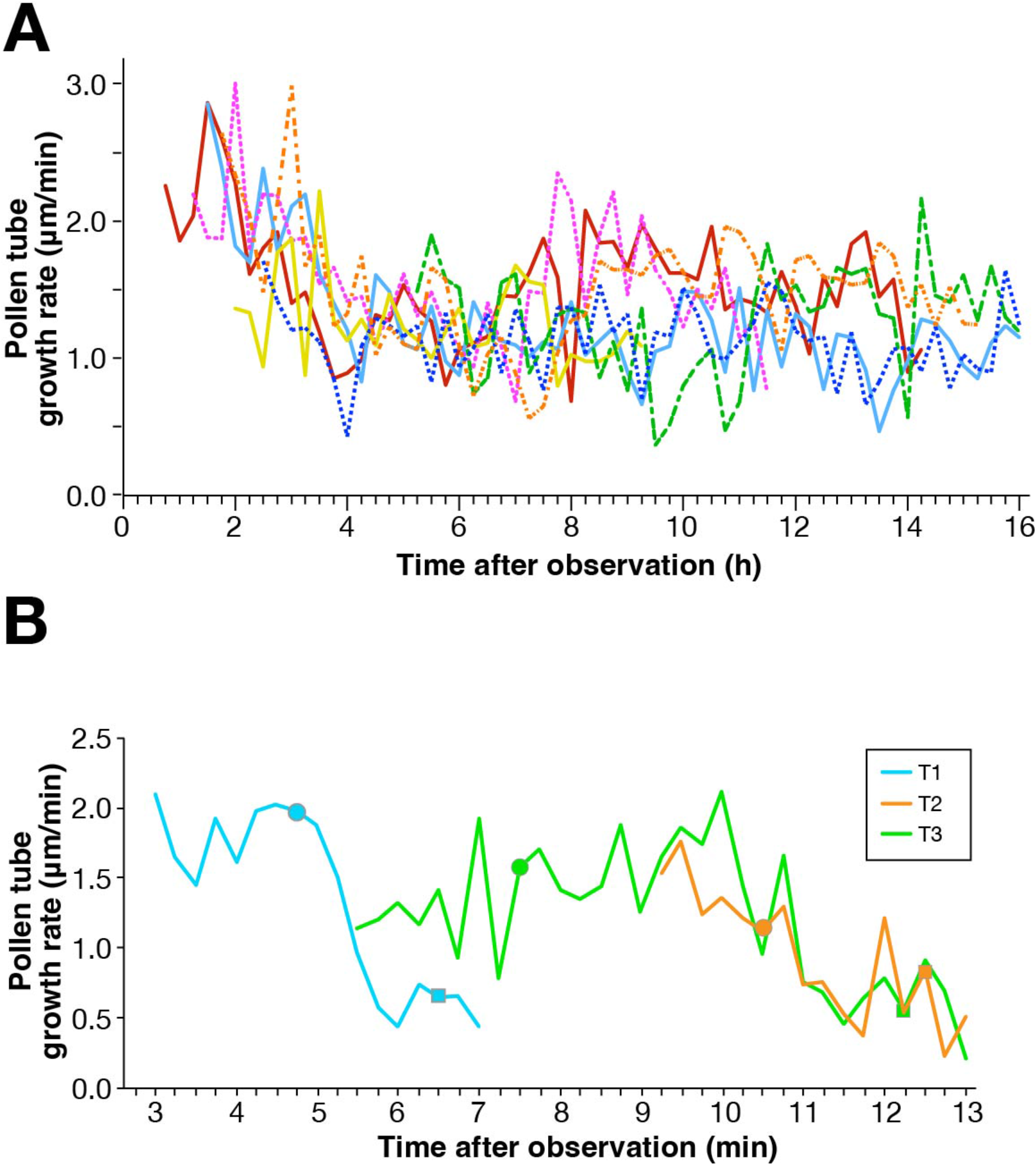

**Fig. S4.**
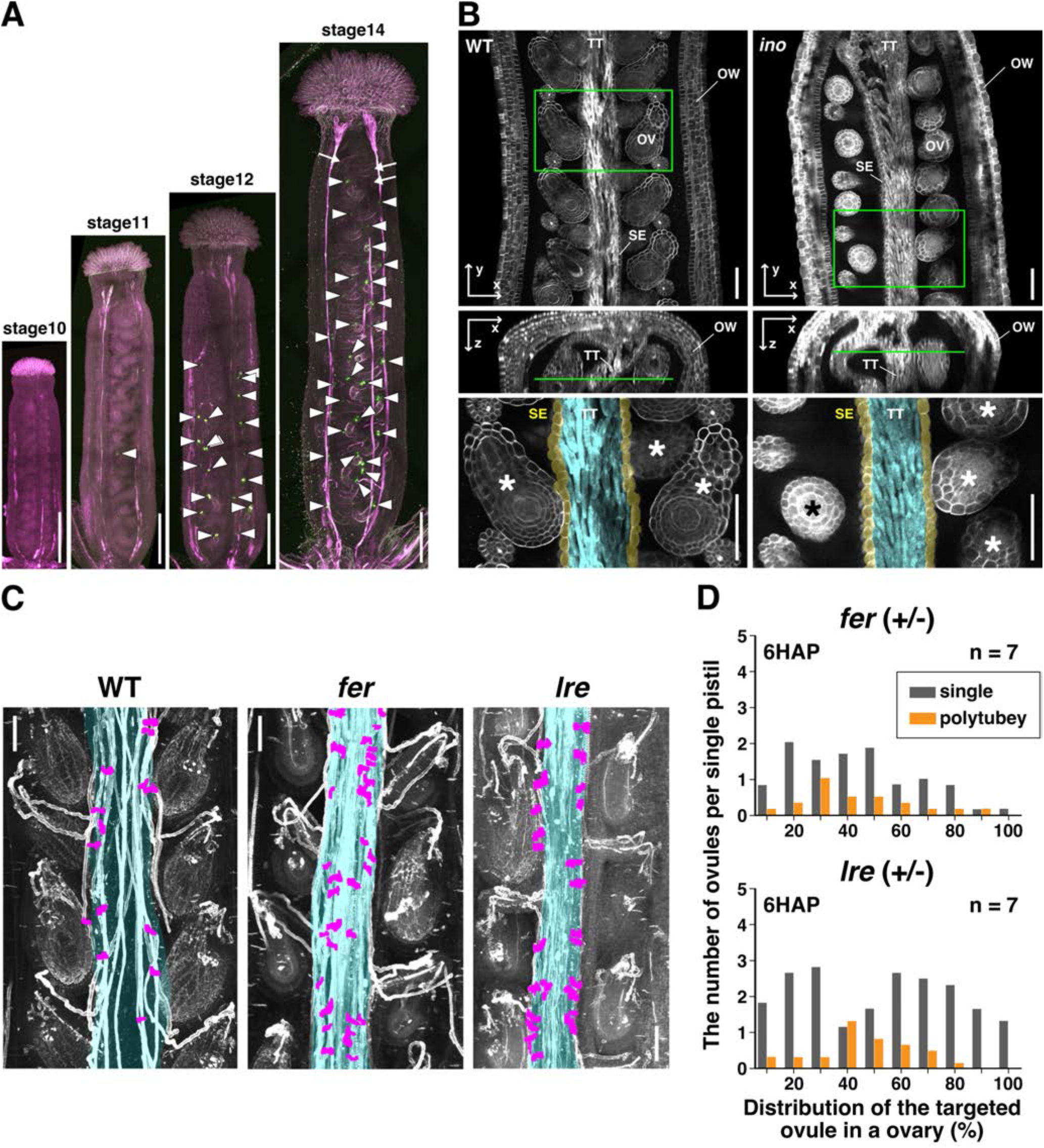

**Table S1.**
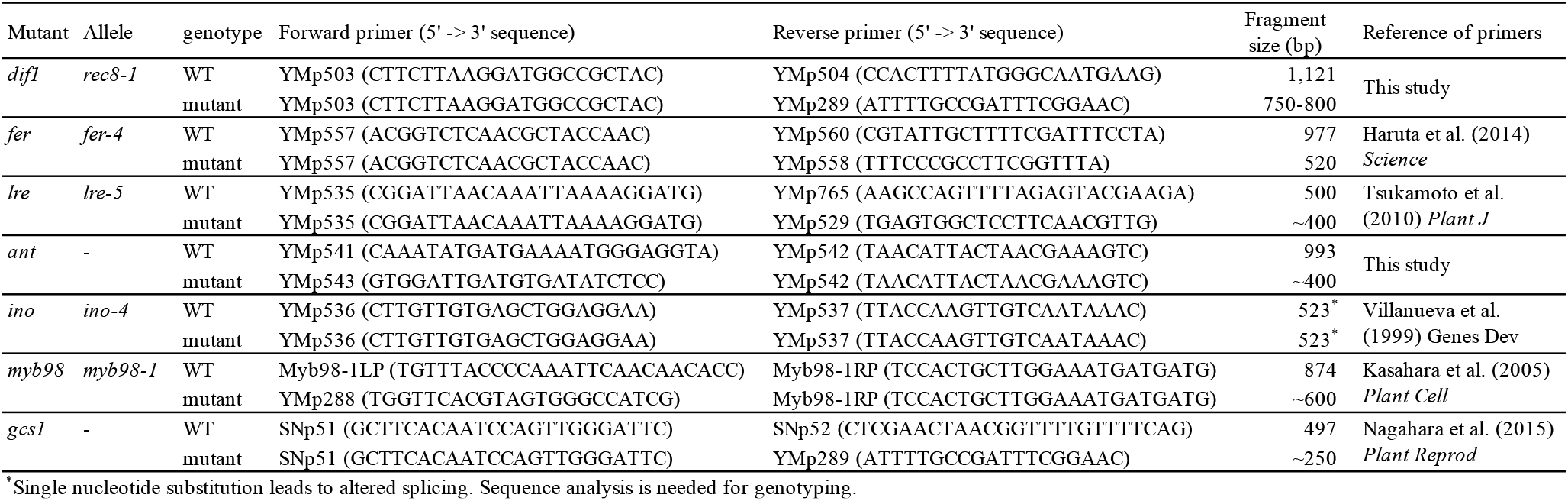
List of primers for mutant genotyping.

