## Supplemental information for "Deep imaging revealed dynamics and signaling in one-to-one pollen tube guidance"

### Materials and Methods

#### Plant materials and growth conditions

*Arabidopsis* ecotype Columbia (*Col-0*) was used as the WT plant. *LAT52p::mTFP1*, *sGFP*, *Venus*, *TagRFP* and *mApple*[1], FGR8.0[2], *MYB98p::GFP*[3], and *HDG11p::NLS-YFP*[4] have been described previously. Nuclei in the septal epidermal cells of *HDG11p::NLS-YFP* are labelled with YFP. Homozygous mutant seeds of *ant* (SALK_022770), *dif1* (*rec8-1*; SALK_091193), *fer* (*fer-4*; GK-106A06), *gcs1* (SALK_135496), *ino* (*ino-4*; N6148), *lre* (*lre-5*; CS66102), and *myb98* (*myb98-1*; SALK_020263), and heterozygous seeds of *fer* (*fer-4*; GK-106A06) and *lre* (*lre-5*; CS66102) were obtained from the Arabidopsis Biological Resource Center at Ohio State University (Columbus, OH, USA) or the GABI-KAT line[5]. Seeds were germinated on agarose plates at 22°C under 24 h light. The primers used for genotyping each mutant are listed in Table S1. Fourteen-day-old seedlings were transferred into a mixture of vermiculite and potting compost. Seedlings were grown at 21–24°C under long-day conditions (16 h light/8 h dark).

**Single-locule method**

The single-locule method was developed for live imaging of pollen tube guidance inside a living pistil (Fig. S2). This method normalizes the optics for *Arabidopsis* two-photon live imaging[1] through the use of a previous method[6]. Fully mature stage 14 flowers were used to avoid potential pistils with immature ovules. The flowers were emasculated 18–24 h before being hand-pollinated. A frame made of a 0.2 mm thick silicone film (AsOne, Osaka, Japan) was placed on the glass bottom dish as a mold (Fig. S2A, Setting), as described previously[7]. Pollen germination medium (PGM)[8] and low-melting point agarose (NuSieve GTG agarose; Lonza Group Ltd., Basel, Switzerland) were poured into the mold. Immediately after hand pollination, a pistil was collected on a glass slide with double-sided tape. Pistils were cut open using an ophthalmic knife (MANI, Inc., Tochigi, Japan) at the valve margins (Fig. S2A, Dissection). The remaining locule was immediately placed horizontally on the cut side (Fig. S2A, Setting), and the flower stalk was embedded in the solid PGM (Fig. S2A, Imaging). Liquid exuding from the solid PGM filled the bottom of the locule by capillary action. A piece of 0.2 mm thick silicone film was then placed on top of the remaining locule as a weight to prevent lifting. The silicone frame was covered with a 0.1 mm thick silicone film to maintain high humidity. Wet paper wipes were placed in a glass-bottom dish to maintain humidity (Fig. S2A, Setting), and the system was sealed with Parafilm (Bemis Flexible Packaging, Oshkosh, WI, USA).

**Two-photon imaging and data analysis**

The imaging system used was based on a previous study[1]. Images were acquired using a laser-scanning inverted microscope (A1R MP; Nikon, Tokyo, Japan) equipped with a 25× water immersion objective lens (CFI 75 Apo 25xW MP, WD = 2 mm; NA = 1.10). A handmade water supply system was equipped with an objective lens, which enabled long-term time-lapse imaging. Emitted fluorescence signals were detected using non-descanned GaAsP PMT detectors (Nikon). To reduce plant autofluorescence and photodamage, excitation wavelengths of 980–1,000 nm were used for live imaging and 850 or 860 nm for imaging of fixed samples[1]. Time-lapse imaging was initiated 1–2 h after pollination (HAP) at 10- or 15-minute intervals. Z-stack images were taken using multiple z-planes at 1–15-μm intervals for 3D construction, projection images, and optical sections inside the ovary. To reconstruct a large image, image stitching was automatically performed using NIS-Elements v4.10 software (Nikon). All images were analyzed with Adobe Photoshop (Adobe Systems, Inc., San Jose, CA, USA) and Fiji [9] software.

**Pollen tube growth rate and pollen tube distribution in the transmitting tract**

Pollen tube growth rates and distribution were calculated by manually tracking the pollen tube tips on the *xy*-projected images using Fiji software. As the pistil itself grew under the single-locule method, an average of three random points on the septum epidermis (SE) were used to normalize the xy-movement associated with pistil development.

**Morphological analysis of the pollen tube and ovule inside the pistil using plant transparent regent ClearSee**

Pistil clearing was performed using the ClearSee.v2 reagent as described previously[10,11]. ClearSee.v2 achieves tissue transparency within the remaining spatial arrangement of organs and fluorescence signals in the *Arabidopsis* pistil[12]. Pistils were collected and fixed with 4% paraformaldehyde (w/v) for 1 h at each time point and then cleared in ClearSee.v2 solution for 4 weeks, according to previous reports[11]. To assess ovule maturation, FGR8.0 immature pistils were fixed before flowering at stages 10, 11, and 12 and then cleared with ClearSee.v2. Pistils at stage 14 were emasculated at stage 12. To observe ovary tissue structure, pistils were stained with 0.1% (w/v) calcofluor staining solution after clearing with ClearSee.v2. Transpared pistils were observed using two-photon excitation microscopy (2PEM) (A1R MP, Nikon) at a suitable excitation wavelength for each fluorescent protein[1].

**Seed distribution under limited pollination**

Stage 12 buds of WT flowers were emasculated and used for experimentation after being allowed to mature for 24 h. For hand-pollination, single pollen grains were applied with tweezers to the center or lateral side of a stigma under a stereomicroscope. After 3 d, 52 pistils with central pollination and 44 pistils with lateral pollination were collected. Sample fixation and clearing by ClearSee.v2 were performed in the same way as described above. After one week of clearing, transparent siliques were observed under a stereomicroscope (SZX7; Olympus, Tokyo, Japan) with an i-NTER LENS optical adapter (Microscope Network Co. Ltd.). The seed-set distribution within the silique was analyzed using Fiji software to determine the percentile of seeds present in each of the ten percentiles of ovary length, from the most apical (1–10%) to the most basal (90–100%) aspect of each ovary.

**Targeted ovule distribution under maximum pollination**

Stage 12 buds of WT flowers were emasculated and used for experimentation after being allowed to mature for 24 h. To achieve maximum pollination, stamens from the opening flower were pinched with tweezers and gently brushed across the emasculated stigma to attach pollen under a stereomicroscope. The 12 pollinated pistils were collected 6 h after pollination. Sample fixation and clearing by ClearSee.v2 were performed as described above. An ovule with at least one pollen tube on its funiculus was defined as the target ovule. Fiji software was used to measure the distribution of the funiculus of the targeted ovule. The percentile of seeds present in each of the ten percentiles of ovary length was determined, from the most apical (1–10%) to the most basal (90–100%) aspect of each ovary. The distribution of 205 ovules and the tips of the most growing pollen tubes in the 12 pistils were analyzed.

**Analysis of pollen tube emergence and polytubey in the pistil using transparency by sodium hydroxide with aniline blue staining**

Pistils were emasculated 18–24 h prior to hand pollination. Pistils from WT, homozygous *fer* (–/–), heterozygous *fer* (+/–), homozygous *lre* (–/–), and heterozygous *lre* (+/–) plants were pollinated with WT pollen. The pollen of the homozygous *gcs1* mutant was pollinated with WT pistils. Pistils collected 4, 6, and 18 h after pollination were fixed with a 9:1 mixture of ethanol:acetic acid and then incubated in 1 N sodium hydroxide (NaOH) for clearing[13]. After overnight incubation, pistils were stained with 0.1% (w/v) aniline blue in $K_3PO_4$ buffer for more than 10 min[14]. The stained pistils were rinsed with sterilized distilled water and then observed using 2PEM at an excitation wavelength of 850 nm (A1R MP, Nikon). The distributions of the ovules showing either single pollen tube attraction or polytubey were analyzed manually with Fiji software. The visualization and statistical analyses of data were performed using GraphPad Prism v9.0 (GraphPad Software, San Diego, CA, USA). To determine the number and location of pollen tube emergence, data were analyzed using one-way analysis of variance (ANOVA), followed by Tukey's post-hoc test.

**Supplementary Figure Legend**

**Figure S1.** Reproductive organ structures and processes of *Arabidopsis thaliana*.

(A) A schematic representation of the structures of the *A. thaliana* flower and the process of pollen tube guidance inside the ovary. A flower has one pistil in the center, which is formed by fusing the carpels. The fused carpels form two locules, which harbor around 20-30 ovules each in a vertical arrangement inside the locule. Merged region forms septum harboring the placentae, from where a stalk-like structure funiculus connects each ovule. When pollen lands on the stigma, pollen germinates pollen tube. Pollen tube penetrates inside the stigma and enters the style connected to the transmitting tract (TT) in the ovary. Pollen tube guidance after entering the TT was divided into three steps in this study: (1) pollen tube emergence from the TT into a locule (pollen tube emergence), (2) pollen tube guidance from the surface of the septum to the funiculus (funicular guidance), (3) pollen tube guidance from the funiculus to the micropyle (micropylar guidance). After micropylar guidance, pollen tube enters the micropyle of the ovule and releases sperm cells in the synergid cells, and then double fertilization with egg cell and central cell occurs. (B–E) Pollen tubes in the TT. WT pistil pollinated with WT pollen collected at 18 hours after pollination was fixed and cleared by 1N sodium hydroxide. Pollen tubes in the pistil were stained with aniline blue. (B) The *xy*-maximum projection image of the ovary in the maximum pollination. Fluorescent signals of pollen tubes and ovary were shown as cyan and magenta. White dotted lines show the ovary wall. (C) Vertical optical section of (B). Pollen tubes inside the TT of the optical *xy* section was shown. (D) Optical cross section of (B) generated by 1-μm steps with 123 planes, which is shown as green line in (B). (D) The number of pollen tube. Pollen tubes in the TT of (C) was counted by multi-point tool in the Fiji software. The 69 pollen tubes in the TT were shown as yellow circle. TT, transmitting tract. Scale bars, 50 μm.

**Figure S2.** The single-locule method.

(A) Schematic representation of sample preparation of the single-loculle method. Pollinated pistils were cut and placed on the mold made with solid pollen germination medium (PGM) in the silicone film at the center of glass bottom dish. One locule was removed without injuring the septum surface to observe pollen tube guidance inside the transmitting tract (Dissection). To maintain high humidity, wet Kimwipes were placed in the glass bottom dish, and sealed with Parafilm (Setting). These procedures must be carried out quickly and under constant temperature control to prevent tissue damage. A pistil placed horizontally on the PGM in a silicone frame for observation by an inverted microscope (Imaging). The bottom side with the locule removed was filled with liquid PGM by capillary action. To keep pistils from moving and drying, silicone films were placed on both the pistil and silicone flame. Two-photon imaging was performed by direction of removed locule. (B) Live imaging of the pollinated pistil by two-photon microscopy under the single-locule method. The *xy*-, *xz*-, and *yz*-projection images were shown. Pistil from *HDG11p::NLS-YFP* was pollinated with mTFP1 expressing pollen. Epidermal nuclei and pollen tube in an ovary were labeled with YFP (magenta) and mTFP1 (green), respectively. The top is the stigma side. Schematic representation was also shown at the bottom. Arrowheads indicate attracted pollen tubes toward the ovule. Arrow indicates the point of pollen tube emergence. (C) The *yz*- and *xz*-optical slice images were shown as white box in B. Fluorescent signals derived from septum epidermis and funicular autofluorescence were shown as yellow and white dotted lines, respectively. PT, pollen tube; OV, ovule; SE, septum epidermis; RL, remaining locule; TT, transmitting tract; FU, funiculus. Scale bars, 100 μm.

**Figure S3.** Pollen tube growth rate in the WT pistils under the single-locule method.

(A) Pollen tube growth rate in the transmitting tract (TT) of the maximum pollinated WT pistil. The pollen tube growth rate of 7 non-emerged pollen tubes in a TT shown in Fig. 1B. Maximum intensity *xy*-projections with images taken at 15-min intervals were used to analyze. See also Movie S1A. (B) The growth rate of 3 emerged pollen tubes in a TT of the WT pistil with limited pollination shown in the Fig. 3B. Maximum intensity *yz*-projections with images taken at 15-min intervals were used to analyze. Filled circles and squares show the time points of the pollen tube attachment to the SE and that of pollen tube emergence, respectively. See also Fig. 3C and Movie S2C.

**Figure S4.** The internal structure of a transparent pistil.

(A) Floral stage dependent ovule development in the WT pistil. Pistils from stage 10 with petals reaching the length of the lateral stamens to the stage 13–14 with opening flower [15] were analyzed. Unpollinated pistils from *FGR8.0* were cleared by ClearSee and observed by 2PEM with 960 nm excitation. Maximum intensity projections for *xy*-projection images were generated from 25–37 z-stack images with 10-µm intervals.

Autofluorescence of pistil was shown in magenta. Arrowheads show the GFP expression in the synergid cells driven by *MYB98* promoter of the *FGR8.0* construct. (B) Cell wall-stained clearing WT and *ino* unpollinated pistils. Optical *xy*- and *xz*-sections were generated by 2-µm steps with 101 planes. Magnified images were shown in the bottom which are shown as green in the upper and middle images. Septum epidermis and TT were colored overlay with yellow and cyan. (C) Pollen tube emerging points in the WT, *fer*, and *lre* mutant ovaries at 24 hours after pollination (HAP). Pollen tubes stained with aniline blue dye and pistil autofluorescence was shown in cyan and gray, respectively. Emerging point on the septum epidermis of each pollen tube is shown as magenta on the *xy*-projection images. (D) The number of ovules showing single pollen tube attraction (gray) and polytubey (magenta) per ovaries of heterozygous fer (+/–) and lre (+/–) at 6HAP. Bar charts represent the number of ovules present in each ten percentile of ovary length from the most apical (0%–10%) to the most basal (90%–100%) of each pistil. Asterisks show the ovule. SE, septum epidermis; TT, transmitting tract; OV, ovule; OW, ovary wall. Scale bars, 200 µm (B), and 50 µm (B and C).

**Movie S1.** Real-time monitoring of pollen tube growth and guidance in the ovary was performed using the single-locule method, as shown in Fig. 1B–G.

This supplemental movie shows time-lapse images of growing pollen tubes and pollen tube guidance in the pistil. (A) WT pistils pollinated with a mixture of pollen from *LAT52p::mTFP1* (cyan) and *LAT52p::TagRFP* (orange) under maximum pollination. Pistil autofluorescence is shown in magenta. Maximum intensity projections for *xy*- (left) and *yz*- (right) projection images were generated from 10 z-stack images with 15 µm intervals using 2PEM with 990 nm excitation. Images were taken at 15 min intervals and the movie was displayed at 10 frames per second. (B and C) Time-lapse images of pollen tube growth and guidance in the pistil of *MYB98p::GFP* pollinated with a mixture of pollen from *LAT52p::mTFP1* (cyan) and *LAT52p::TagRFP* (orange). Synergid cells in the ovule were labelled with GFP (yellow). Maximum intensity projections for *xy*- (left) and *yz*- (right) projection images were generated from 12 z-stack images with 15 µm intervals using 2PEM with 990 nm excitation. Images were taken at 10 min intervals and the movie was displayed at 10 frames per second. (C) Funicular guidance of a single pollen tube. The numbers stamped in each frame indicate the time (h:mm) from the start of observation. Asterisks and T indicate synergistic cells and targeted synergistic cells expressing GFP, respectively. Arrows and arrowheads indicate the tips of the fastest growing and attracted pollen tubes, respectively. Scale bars, 100 µm.

**Movie S2.** Attracted and non-attracted pollen tubes in the pistil under limited pollination, as shown in Fig. 2G–J and Fig. 3B, respectively.

This supplemental movie presents time-lapse imagery of pollen tube guidance in the pistil of *HDG11p::NLS-YFP* (magenta) pollinated with *LAT52p::mTFP1* (green) pollen. Maximum intensity projections for *xy*- (left) and *yz*- (right) projection images were generated from 18 (A and B) or 17 (C) z-stack images with 10 µm intervals using 2PEM with 980 nm excitation. Images were taken at 15 min intervals and the movie was displayed at 15 frames per second. The numbers stamped in each frame indicate time (h:mm) from the start of observation. Asterisks and T indicate non-targeted and targeted ovule autofluorescence, respectively. Arrow: attracted pollen tube; arrowhead: non-attracted pollen tubes. Scale bars, 100 µm.

**Movie S3.** Polytubey in WT, *fer*, and *lre* pistils (Fig. 6A–C).

This supplemental movie shows time-lapse images of pollen tube guidance and polytubey in the ovary. (A) WT pistil expressing *HDG11p::NLS-Venus* (magenta) pollinated by *LAT52p::TagRFP* (green) pollen. Maximum intensity projections for *xy*- (left) and *yz*- (right) projection images were generated from 21 z-stack images with 7 µm intervals using 2PEM with 990 nm excitation. Images were taken at 15 min intervals and the movie was displayed at 8 frames per second. The numbers stamped in each frame indicate time (h:mm) from the start of observation. Asterisks indicate the attracted single pollen tubes. Polytubey ovules indicated with T. Two pollen tubes attracted to the same ovule are pointed out with arrows. (B and C) *fer* (A) and *lre* (C) pistils pollinated with *LAT52p::TagRFP* (green) pollen grains. Maximum intensity projections for *yz*- (right) projection images were generated from 20 (A) or 23 (C) z-stack images with 8 µm intervals using 2PEM with 1,000 nm excitation. Images were taken at 15 min intervals and the movie was displayed at 8 frames per second. The numbers stamped in each frame indicate the time (h:mm) from the start of observation. Pollen tubes attracted to the same ovule are shown as arrowheads. Pollen tubes that turn back to the septum are indicated with arrows. Scale bars, 100 µm.

### AUTHOR CONTRIBUTIONS

Y.M. designed the experiments and directed the project. Two-photon imaging using the single-locule method was developed by Y.M. The handmade water-supply system for two-photon time-lapse imaging and the ClearSee method were developed by D.K. The method to analyze the number and distribution of pollen tubes under aniline-blue staining was established by T.T.N. and performed by D.S and I.K. Other experiments were performed by Y.M. and S.N. The manuscript was written by Y.M., with comments from D.S., S.N., T.T.N., D.K., and T.H.

### ACKNOWLEDGMENTS

We thank Dr. F. Berger and Dr. U. Grossniklaus for their helpful suggestions and discussion. We thank Dr. R. Groß-Hardt, Dr. R. Palanivelu, Dr. R.D. Kasahara, Dr. M. Ueda, Dr. D. Maruyama, and Dr. D. Susaki for providing plant materials and plasmids, Dr. H. Takeuchi and Dr. Y. Hamamura for providing plasmids, and S. Nasu, T. Nishii, and T. Shinagawa for assistance in preparing the materials. Microscopy was conducted at the WPI-ITbM of Nagoya University and supported by the Advanced Bioimaging

Support through MEXT/JSPS KAKENHI (22H04926). This work was supported by grants from the Japan Science and Technology Agency (ERATO Grant number JPMJER1004 to T.H. and FOREST Program Grant Number JPMJFR204T to D.K.); the Japan Society for the Promotion of Science: Grant-in-Aid for Transformative Research Areas (20H05778, 20H05779 to Y.M.)]; the Ministry of Education, Culture, Sports, Science and Technology in Japan (no. 18K14741 to Y.M., no. JP22H04668 to D.K., and no. JP16H06465 to T.H.); and the Program for Promoting the Enhancement of Research Universities (2022 to Y.M.).
